# Parental feeding preferences rather than sibling competition determine the death of smaller nestlings in asynchronous broods

**DOI:** 10.1101/2020.12.01.404590

**Authors:** Manuel Soler, Francisco Ruiz-Raya, Lucía Sánchez-Pérez, Juan Diego Ibáñez-Álamo

## Abstract

Hatching asynchrony is a reproductive tactic that, through the creation of competitive hierarchies among offspring, allows parents for a quick adjustment of brood size via the death of smaller nestlings. This strategy is considered to be adaptive in case of unpredictable and/or poor environments in which it would guarantee that at least larger nestlings will fledge. Brood reduction is the usual outcome in asynchronously hatched broods since first-hatched nestlings are larger and get a disproportionately larger share of the food delivered by parents, often leading the youngest nestling to starve to death soon after hatching. However, we still do not know the proximate mechanisms of such brood reduction. One possibility is that the smallest nestling is not fed because larger nestlings outcompete it, which implies that nestlings control resource allocation. Alternatively, parents might actively ignore the persistent begging from their smallest nestling, which would involve that parents control food allocation. To determine whether parents or nestlings ultimately induce brood reduction in this situation, we experimentally created asynchronous broods of Eurasian blackbird (*Turdus merula*) nestlings and quantified food allocation by parents in two different situations: when sibling competition was allowed and, alternatively, when competition was prevented by physically separating nestlings within the nests by using wooden barriers. Our results showed that experimentally introduced smaller nestlings received less food than their larger nestmates both when competition among nestlings was allowed and when it was prevented. When adult males and females are considered separately, males fed the smallest nestling less often regardless of whether sibling competition was allowed or not, but adult females showed no differences. We can conclude that the smallest nestling starves mainly because parents actively ignore its begging. The higher competitive ability of the larger nestlings seem to have little effect given that although the smallest nestling is fed at a higher rate when physical interactions are prevented by the wooden barrier than when not, this difference is not significant. These findings suggest that parents rather than nestlings have the main control over food allocation.

## Introduction

In birds, the apparently most effective way to maximize parental fitness is to ensure that all laid eggs eventually result in fledglings. This is indeed what happens in most species: parents start incubation after clutch completion producing a brood of synchronously hatched nestlings of similar size. This strategy avoids the existence of competitive hierarchies among offspring and favours an equalitarian allocation of food among nestlings. Consequently, parents increase the chances that all nestlings will survive to fledge by preferentially feeding the hungriest nestling according to signal of need models (Davis et al. 1999; Soler 2001; Jeon 2008; Caro et al. 2008). Based on available information, it is often assumed that both hatching synchrony and food allocation favouring nestlings in worse condition (i.e. those begging at higher intensity) provide adaptive benefits in predictable and good environments since, when food is abundant, a high provisioning rate would allow parents to provide more food to the undernourished nestling and all offspring will be able to fledge (Lack 1947, 1954; Magrath 1990; Davis et al. 1999; Jeon 2008; Caro et al. 2016). In contrast, a synchronous brood could compromise nestling survival in unpredictable and/or poor environments since feeding all nestlings when food is scarce could provoke starvation in all or most of them. Thus, when food is scarce, any tactic allowing parents to adjust brood size according to food availability would be adaptive because this would guarantee that at least larger nestlings will fledge (Lack 1954; Jeon 2008; Caro et al. 2016). Hatching asynchrony is considered one of such tactics since, in unpredictable and/or poor environments, a size hierarchy among offspring would allow for a quick adaptive adjustment of brood size through death of smaller nestlings (the brood reduction hypothesis (Lack 1945, 1954)).

Brood reduction is the usual outcome in asynchronously hatched broods, in which first-hatched nestlings are larger and exhibit a higher competitive ability that allow them to outcompete later-hatched (smaller) nestlings by getting a disproportionately larger share of the food delivered by parents (Lack 1954; Soler 2001; Jeon 2008). As a consequence, the youngest nestling often starves soon after hatching. This is supported by the fact that a bigger size increases the ability of a nestling to jostle for the best position in the nest, which is considered crucial to successfully compete for food (McRae et al. 1993; Malacarne et al. 1994; Kilner 1995; Ostreiher 2001). Preferential feeding of larger nestlings has usually been interpreted as the outcome of scramble competition among offspring, assuming that nestlings exert full control over resource allocation by parents (scramble competition models (Macnair and Parker 1979; Mock and Parker 1997; Parker et al. 2002)). However, empirical and experimental evidence has shown that larger nestlings obtain more food than smaller nestlings even when showing a lower begging behaviour (Smiseth and Amundsen 2002: Mock et al 2011), which indicates that parents may have at least partial control in food allocation within asynchronous broods (Krebs et al 1999; Smiseth et al 2003). Despite this evidence, to what extend parents and nestlings exert control of brood reduction still remains unclear (McRae et al. 1993; Mock et al. 2009). Recent studies suggest that parental control is much more relevant than nestling behaviour. For example, house wrens (*Troglodytes aedon*) are able to switch from synchronous hatching with all nestlings surviving to fledge in favourable conditions, to asynchronous hatching with brood reduction in less favourable conditions, even within the same breeding season (Ellis et al 2001). In Eurasian hoopoes (*Upupa epops*), it has been reported that while males feed larger nestlings placed in the best position (entrance of the nest), females usually enter the nest cavity and feed preferentially the smallest nestlings (Ryser et al. 2016). This seems to be the case in many asynchronously hatching species, in which females (and not males) feed the smallest nestling of the brood showing that females have control over food distribution (Gottlander 1987; Stamps et al. 1987; Lifjeld et al. 1992; Leonard and Horn 1996; Slagsvold 1997a, 1997b; Krebs et al. 1999; Ploger and Madeiros 2004; Dickens and Hartley 2007; Budden and Beissinger 2009; Lahaye et al. 2015; Ryser et al. 2016). The frequently reported fact that females (and not males) feed the smallest nestling in the brood has inspired the “Male manipulation hypothesis”, which states that females, by provoking hatching asynchrony and keeping alive the smallest nestling (runt), which begs more and at a higher intensity, force males to work harder, carrying more food to the nest in asynchronously hatched broods (Soler et al. In prep.).

The fact that males and females differ in their food allocation rules in asynchronously-hatched broods implies that at least one of the sexes (males that are feeding selectively the best competitors, or females that are feeding selectively smaller nestlings) allocates food ignoring nestling begging behaviour, thus exerting the main control over food allocation. The crucial point in this situation is the undernourished smallest nestling who, after becoming a runt (i.e. a nestling with a retarded growth that does not have any probability of survival), starve to death. There are three possibilities to explain why one or more smaller nestlings in asynchronous broods starve: (a) runts are not fed because of sibling competition (i.e. larger nestlings outcompete them), (b) parents actively ignore the persistent begging from their smaller nestling, or (c) a mix of both. As far as we know, no study has assessed the relative importance of these three possibilities in the context of brood reduction. In this experimental study, we aim to determine to what extend either sibling competition or parental preferences are the responsible of runt’s starvation. Since previous studies have shown that males and females can follow different rules when allocating food among offspring (see references above), we considered the role of both sexes separately. To fulfill these objectives, we experimentally created asynchronous nests to explore parental food allocation patterns in two different situations: (i) allowing physical interactions among nestlings and (ii) impeding them by using a wooden barrier that forces each nestling to stay in its portion of the nest.

Our main hypothesis, considering the increasing evidence that control by parents in food allocation is more relevant than previously thought (see references above), is that parents actively discriminate against runt nestlings regardless of nestling competition and nestling need (Hypothesis 1). This hypothesis predicts that the smallest nestling will receive less food than its nestmates not only when physical interactions are allowed, but also when the presence of a wooden barrier would impede those interactions. The other two non-mutually-exclusive alternative hypotheses are: food allocation by parents is determined by scramble competition among nestlings (Hypothesis 2). This hypothesis predicts that in natural conditions, when physical interactions occurs, the smallest nestling will receive less food than its nestmates, but when the presence of a wooden barrier would impede larger nestlings blocking their smaller nestlings, the neediest nestling, should be fed preferentially, or at least at the same frequency as its larger nestmates. Finally, the intermediate possibility is that both adults and offspring exert some control over food allocation by parents (Hypothesis 3). This hypothesis predicts that the smallest nestling should be fed at a significantly higher rate when physical interactions are prevented by the wooden barrier than when not.

## Methods

### Study area and species

Fieldwork was carried out in the Valley of Lecrín (Southern Spain; 36° 56′ N, 3° 33′ W; 580 m a.s.l.) between mid-March and June in 2013 and 2015. As model species, we used the Eurasian blackbird (*Turdus merula*, hereafter blackbird), a medium sized passerine with a clear sexual dimorphism (adult males being black with a distinctive yellow-orange beak and eye-ring, while adult females are dull dark, with lighter brown streaks on their breast), which makes it easy to recognize the sex involved in each feeding event. In our study area blackbirds’ clutch size ranges from 2 to 5 eggs (Pers. Observ.). We have chosen this species because females start incubation before the last egg is laid inducing moderate hatching asynchrony (the last egg hatches latter than the rest and in some cases brood reduction occurs, mainly at the end of the breeding season and in other situations of poor environmental conditions (Cramp 1985)).

### General field procedures and experimental design

We actively searched for blackbird nests in the study area throughout the breeding season. Once located, nests were visited every two days and, close to hatching, daily in order to detect newly-hatched chicks. In this species, hatching order can easily be assessed by a daily nest checking; even so, in those cases in which two nestlings hatched within 24 hours, we relied on the chick’s weight to establish the nestling rank. Recently-hatched nestlings were marked on their tarsus by using coloured permanent markers (Lumocolor) and, on days 6-7 post-hatching, with numbered rings for individual recognition.

We experimentally created four-nestlings asynchronous broods by conducting a cross-fostering manipulation in nests with similar hatching dates. On the experimental day 0, a one-day old nestling from a donor nest (experimental nestling; mean weight = 7.4 ± 0.3 g, n = 14) was introduced into a recipient nest containing three-days old nestlings (± 1 day, siblings’ mean weight = 14.5 ± 0.3 g, n = 42). The next day (experimental day 1; 9:00 – 13:00 h), parental feeding activity was video recorded in two consecutive and non-overlapping trials, each trial lasting 1.5 hours. In one of these trials, siblings were allowed to physically compete for food while, in the other trial, physical interaction among nestlings was prevented by placing a wooden cross-shaped barrier into the cup nest (attached to the base of the nest by a twine thread) so that siblings remained separated in four same-size compartments. Barriers were placed 30 minutes before the start of the recordings so that parents could get used to their presence in the nest. Nestlings were randomly placed into the nest compartments, being able to raise their heads above barriers during begging. In all cases, parents could easily feed all nestlings from the edge of the nest. The experimental trial in which nestlings remained separated by the wooden barrier was alternated in successive nests (first trial or second trial) to clearly separate the effect of the barrier presence from the potential order effect. We avoided altering the nest structure (i.e. relative position of nestlings in the nest) in successive trials. Although previous studies have suggested that the position of a nestling in the nest relative to its nestmates and the feeding parent may influence food obtained by the nestling (McRae et al. 1993; Malacarne et al. 1994; Kilner 1995; Ostreiher 2001), we assume that positional effects are irrelevant in our study because of two reasons. First, barriers and nestlings were placed randomly with regards to the position of the parent, and second, our barrier separating the nest cup in four parts removes the central position, which may be the most favourable.

About 30 minutes before the first trial, a video camera (Panasonic HDC-SD40) was placed near the nest (3 - 5 m) to determine parental feeding decisions and quantify food delivery rates to the first-hatched nestling (a-nestling), the second-hatched nestling (b-nestling), the third-hatched nestling (c-nestling) and the smallest (experimental) nestling. The video camera was attached to a small tripod and hidden in the vegetation near the nest, often at a higher height than the nest so that the filming was made from above in most cases (angles between 15° - 45°). Food allocation patterns were investigated by assessing the overall food load received by each nestling in one hour. To do that, we quantified the number of food items delivered to each chick by parents. One food item is defined as a prey that represents a volume similar to the size of blackbirds’ bill. Thus, the size category of food items was estimated from recordings by comparing the volume of each food item relative to the bill volume of the parent within bins of 50% (e.g., 50%, 100%, 150%, etc., of bill volume) (Hauber and Moskát 2008). Additionally, food items smaller than 50% of the bill volume were all quantified as 10% of bill volume as they were mostly small insects of similar size. Finally, the volumes (i.e. size categories) of all items delivered to a single nestling were summed to generate the variable “food load”. In order to distinguish different nestlings in recordings, we marked their bill with individual colours by using permanent markers.

### Statistical analysis

To assess the effects of our experimental manipulation on parents’ food allocation patterns, we fitted linear mixed models (LMM) by using the *nlme* package (Pinheiro et al. 2014), with the number of food items that nestlings received per hour (log-transformed) as a dependent variable. As fixed factors, our models included barrier presence (yes/no), nestling rank (a/b/c/experimental) and the interaction between these two terms, while chick identity nested within nest identity was included as random terms. Only significant interactions were retained in the models (Engqvist 2005). Post hoc contrasts were performed using the *lsmeans* package (Lenth 2016). The proportion of variance explained (R^2^) for our mixed models was calculated according to Nakagawa and Schielzeth (2013) and Nakagawa et al. (2017). In short, two values of R^2^ were calculated: the marginal r-squared (R^2^LMM(m)), describing the proportion of variance explained by fixed factors alone, and the conditional r-squared (R^2^LMM(c)), describing the proportion of variance explained by both fixed and random terms. Assumptions of normality and homogeneity of variances were verified through the visual inspection of residual graphs. All analyses and graphs were performed using R version 3.6.1 (R Core Team 2019).

## Results

### Food allocation: general patterns

The overall amount of food carried to the nest was similar when nestlings were separated by wooden barriers or when they were free to compete for resources (F_1,52_ = 0.24; *p* = 0.62; Fig. 1). Importantly, the food was unevenly distributed among nestlings (F_3,52_ = 9.58; *p* < 0.0001; Fig. 1), regardless of whether physical sibling competition was allowed or not, as indicated by the non-significant two-way interaction (LMM: nestling rank x barrier presence: F_3,49_ = 0.83; *p* = 0.486). These results are in agreement with the prediction derived from Hypothesis 1, while contradict that derived from Hypothesis 2. Overall, the younger experimental nestling received less food than the a-nestling, the b-nestling and the c-nestling (Table 1a; Fig. 1). Regarding to the prediction from Hypothesis 3, although the smallest experimental nestling received more food items per hour (1.34 ± 0.26; n = 14) when separated by the wooden barrier than in natural conditions (0.80 ± 0.18 food items per hour, n = 14), this difference was not significant (estimate ± se = 0.228 ± 0.144; df = 49; t = 1.59; *p* = 0.12). Our statistical model explained 47.6% of total variance, in which the fixed part (i.e. the additive effects of sibling competition and parental preferences) explained 19.7% (Table 1a); however, whether sibling competition is excluded from models, the fixed part still explains 19.7% of variance, which seems to confirm that, while scramble competition among nestlings slightly (although not significantly) reduced the amount of food received by the smallest chick (Fig. 1), food allocation was mainly determined by parental decisions.

**Table 1.** Summary of linear mixed models (LMM) fitted to assess the effect of nestling rank and barrier presence on parental food allocation. Non-significant interactions were not retained in the models. Table shows log-transformed estimates.

| **a) Food allocation: general patterns** |  |  |  |
| --- | --- | --- | --- |
| | $\beta$ (se) | t-value | p-value |
| Intercept | 0.662 (0.072) | 6.142 | 0.000 |
| c-nestling | 0.335 (0.120) | 2.801 | **0.007** |
| b-nestling | 0.638 (0.121) | 5.284 | **< 0.0001** |
| a-nestling | 0.417 (0.121) | 3.456 | **0.001** |
| barrier presence (yes) | -0.036 (0.072) | -0.494 | 0.620 |
| *Random effects* |  | *Variance* |  |
| Nest ID |  | 0.206 |  |
| Chick ID *within* Nest ID |  | 0.181 |  |
| Residual |  | 0.376 |  |
| $R^2_{LMM(m)} = 0.197$ | | | |
| $R^2_{LMM(c)} = 0.476$ | | | |
| **b) Food allocation by males** |  |  |  |
| | $\beta$ (se) | t-value | p-value |
| Intercept | 0.511 (0.102) | 5.030 | 0.000 |
| c-nestling | 0.171 (0.115) | 1.481 | 0.144 |
| b-nestling | 0.520 (0.115) | 4.520 | **< 0.0001** |
| a-nestling | 0.391 (0.115) | 3.392 | **0.001** |
| barrier presence (yes) | -0.067 (0.073) | -0.920 | 0.362 |
| *Random effects* |  | *Variance* |  |
| Nest ID |  | 0.182 |  |
| Chick ID *within* Nest ID |  | 0.144 |  |
| Residual |  | 0.384 |  |
| $R^2_{LMM(m)} = 0.171$ | | | |
| $R^2_{LMM(c)} = 0.393$ | | | |
| **c) Food allocation by females** |  |  |  |
| | $\beta$ (se) | t-value | p-value |
| Intercept | 0.229 (0.090) | 2.552 | 0.014 |
| c-nestling | 0.253 (0.104) | 2.433 | **0.018** |
| b-nestling | 0.638 (0.104) | 2.900 | **0.005** |
| a-nestling | 0.417 (0.104) | 1.200 | 0.235 |
| barrier presence (yes) | 0.059 (0.066) | 0.884 | 0.381 |
| *Random effects* |  | *Variance* |  |
| Nest ID |  | 0.148 |  |
| Chick ID *within* Nest ID |  | 0.126 |  |
| Residual |  | 0.349 |  |
| $R^2_{LMM(m)} = 0.085$ | | | |
| $R^2_{LMM(c)} = 0.301$ | | | |

**Figure 1.**
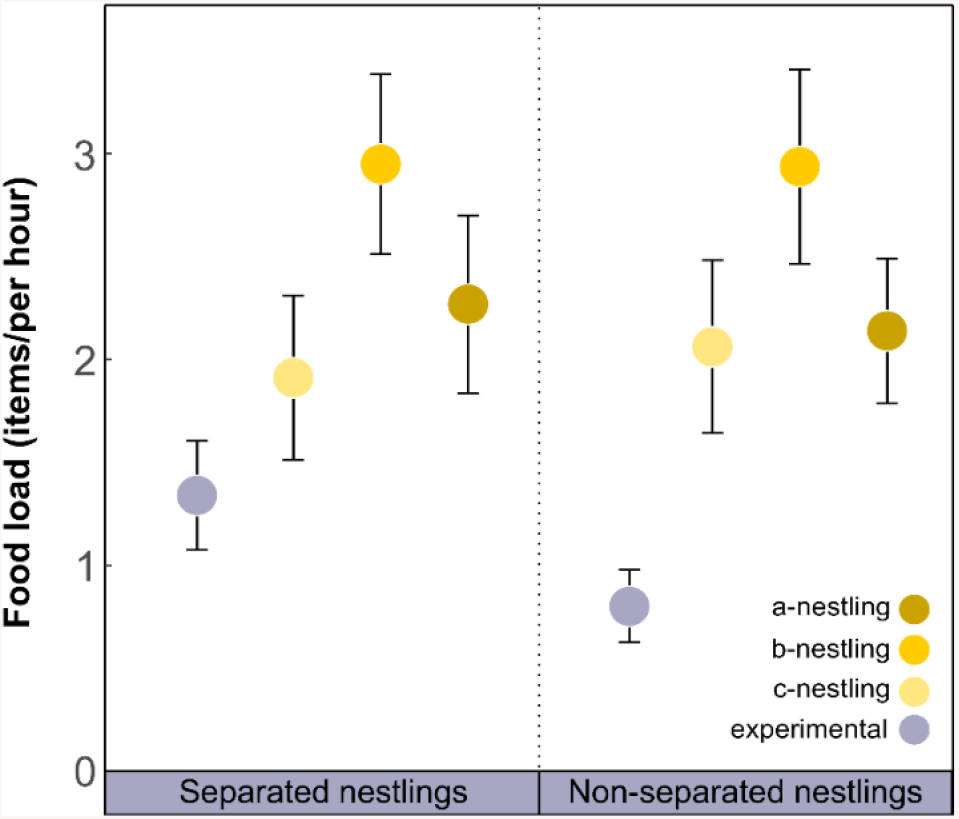
Parental food allocation among non-experimental siblings (n = 42; yellow palette) and the smaller experimental nestling (n = 14; grey). Values are presented as means ± se.

### Food allocation by males

On average, male blackbirds visited their nests to feed nestlings 4.01 ± 0.39 times per hour (range = 1.94 - 7.03, n = 14). Males did not vary the overall amount of food provided to nestlings when they were separated and unable to compete compared with the control situation (F_1,52_ = 0.80; *p* = 0.38). However, males differentially distributed food among siblings (F_3,52_ = 8.05; *p* = 0.0002), providing less food to the smallest nestling regardless the competitive context in which nestlings were maintained (LMM: nestling rank x barrier presence: F_3,49_ = 0.55; *p* = 0.648; Figure 2A). These results support the prediction derived from Hypothesis 1, while contradicts the prediction associated with Hypothesis 2. More specifically, the smallest (experimental) nestling received significantly less food than the a-nestling and the b-nestling, but there were no significant differences between the smallest and the c-nestling (Table 1b; Figure 2A). With respect to the prediction from Hypothesis 3, although the smallest nestling received more food items per hour when separated by the wooden barrier (0.86 ± 0.21; n = 14) than in natural conditions (0.58 ±0.12 feeds per hour; n = 14), this difference was not significant (estimate ± se = 0.112 ± 0.147; df = 49; t = 1.20; *p* = 0.49). Our statistical model for males explained 39.3% of total variance, in which the fixed part explained 17.1% (Table 1b); however, whether sibling competition is excluded from models, parental preferences still explain 16.8% of variance. Taken together these results indicate that, in the case of males, food allocation was mainly determined by parental decisions.

**Figure 2.**
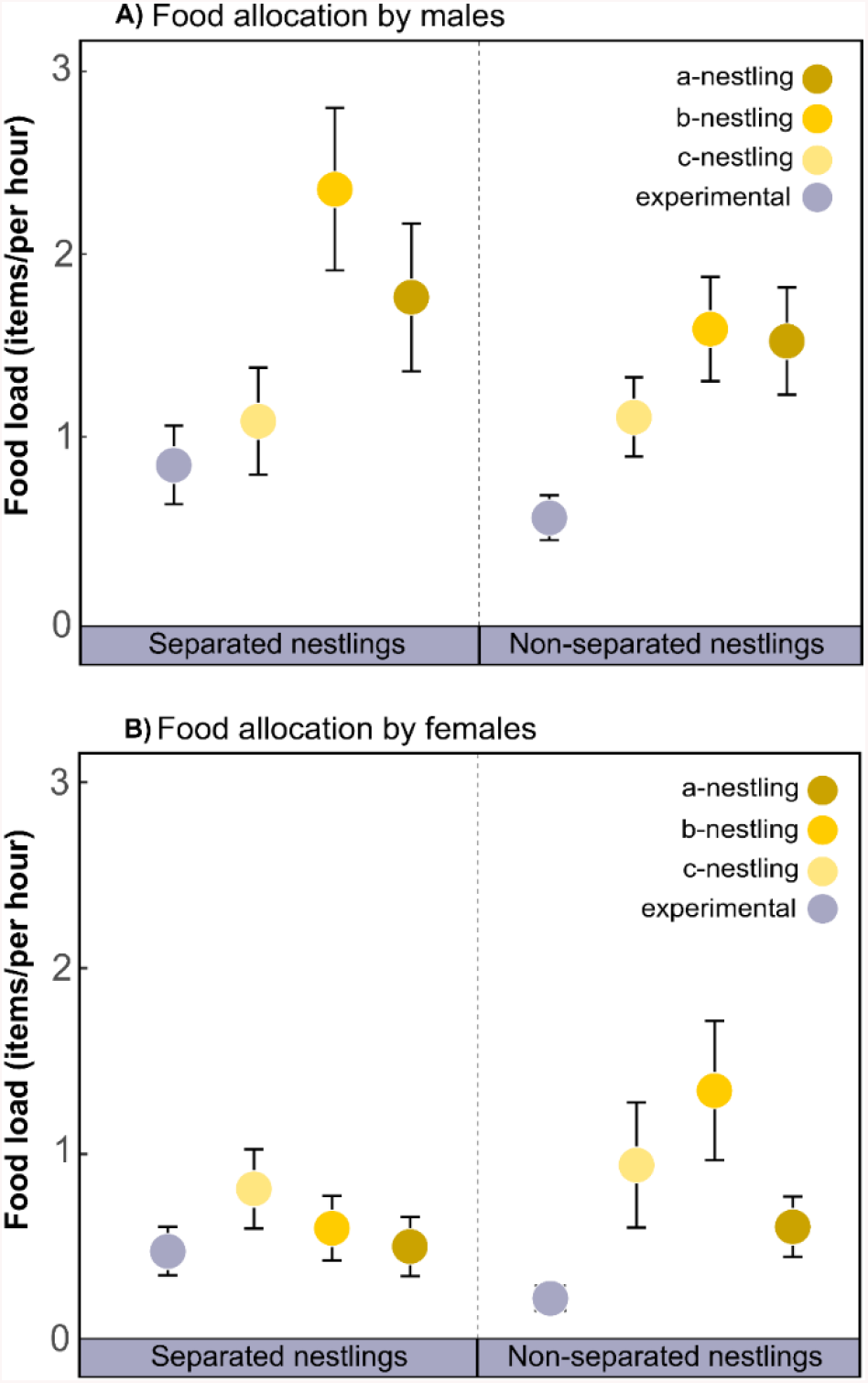
Food allocation among non-experimental siblings (n = 42; yellow palette) and the smaller experimental nestling (n = 42; grey) by male (A) and female blackbirds (B). Values are presented as means ± se.

### Food allocation by females

Our results showed that female blackbirds visited their nests to feed nestlings 2.27± 0.29 times per hour (range = 0.66 – 4.23, n = 14). The overall food load that nestlings received from females were not affected by the absence of physical sibling competition (F_1,52_ = 0.81; *p* = 0.37). Females differentially fed nestlings (F_3,52_ = 3.40; *p* = 0.02), although such differences tended to soften when nestlings could not compete for food (LMM: nestling rank x barrier presence: F_3,49_ = 2.27; *p* = 0.092; Fig. 2B). Interestingly, post hoc tests revealed that, in the absence of physical competition (i.e. nestlings separated by barriers), the smallest experimental nestling was fed by females at similar rates than the other three nestlings (all cases *p* > 0.22). However, when nestlings were free to compete, the smallest nestling received less food than the b-nestling (estimate ± se = −0.533± 0.137; df = 49; t = −3.884; *p* = 0.0003) and the c-nestling (estimate ± se = −0.332± 0.137; df = 49; t = − 2.420; *p* = 0.019), and tended to receive less food than the a-nestling (estimate ± se = - 0.240± 0.137; df = 49; t = −1.752; *p* = 0.086). Taken together, these results support the Hypothesis 2 stating that food allocation by females is mainly determined by nestling competition. However, the fact that the experimental smallest nestling did not receive significantly more food when scramble competition by larger nestlings was prevented seems to support the Hypothesis 1 too, advocating for a key role of parental decisions. Specifically, although the smallest experimental nestling received more food items per hour when separated by the wooden barrier (0.48 ± 0.13; n = 14) than in natural conditions (0.22 ± 0.06 food items per hour; n = 14), this difference was again not significant (estimate ± se = 0.154 ± 0.128; df = 49; t = 1.20; *p* = 0.24). Our statistical model explained 30.1% of total variance, in which the fixed part explained 8.5% (Table 1c). Nevertheless, when sibling competition is excluded from models, the fixed part (i.e. parental decisions) still explains 8.5% of variance.

### Survival of nestlings in asynchronous broods

By day 6 post-hatching, a large proportion of experimental nestlings had starved in experimentally asynchronous broods (42.9%; N = 14). Furthermore, 50% of experimental nests were predated before fledgling, so eventually only one experimental nestling successfully fledged (7.1%).

## Discussion

In species with asynchronously hatched broods, the smallest nestling obtains less food than their larger nestmates and frequently dies by starvation soon after hatching (Lack 1945, 1954; Jeon 2008; Caro et al. 2016). Our results match these previous findings indicating parental favoritism towards larger nestlings. In blackbirds, experimentally introduced smaller nestling received less food than their three larger nestmates (Fig. 1), which usually resulted in the death of the former by starvation. Regarding the main aim of our experimental study, we can state that larger nestlings are not preferentially fed by their parents because of their higher competitive abilities, given that differences in food allocation were maintained in the absence of sibling competition (Fig. 1). The experimental smallest nestling did not receive significantly more food from their parents in the absence of scramble competition than when physical interactions among nestlings were allowed (Fig. 1). Even though a small increase in the food received per hour by the smallest nestling occurs when nestlings are separated by the barrier (Fig. 1), it still receives less food from their parents compared to their lager nestmates. Taken together, our results do not support Hypotheses 2 and 3. However, we have found strong support for Hypothesis 1, which states that food allocation is mainly determined by parental decisions rather than sibling competition. These results imply that, at least in this passerine species with moderate brood reduction, parents have the main control over resources allocation. This could be also the case in other groups such as ardeids, in which brood reduction is much more frequent and larger nestlings even attack smaller ones without parental interference (Mock and Parker 1997). Ploger and Madeiros (2004), in an experiment performed using plastic barriers in the great egret (*Ardea alba*), found that while in natural conditions the largest nestling obtain more food than the other two nestlings in the brood, when nestlings were separated preventing physical interactions, the second nestling in the size hierarchy received more food than the first and the third nestlings. This means that also, in this species, parents may have some control. In our experimental design we have not controlled for nestling position in the nest cup with respect to the feeding parent. However, we are confident that positional effects are irrelevant in our study because barriers and nestlings were placed randomly. Furthermore, several experimental studies in which the relative nestling position was controlled for have found that parents feed their nestlings regardless of their position in the nest (e.g. Kilner 1995, Tanner et al. 2008, Smith et al. 2017).

Interestingly, we found differences between sexes in food allocation. In our study, blackbird males provided significantly less food to the smallest nestling regardless the competitive context in which they were maintained (Fig 2a), which means that males do not simply feed the dominant nestling. When physical competition is prevented, males also preferentially feed larger nestlings. Importantly, this implies that nestlings have little or no control over food allocation regarding males’ feeding events. On the other hand, females did not provide significantly less food to the smallest nestling in the barrier situation, but instead they evenly distributed the food among nestlings (Fig 2b). These results are similar to those found in the great egret. In their study, Ploger and Madeiros (2004) showed that when nestlings were prevented from aggressive interactions, females, but not males, did not feed preferentially the largest nestling, but the second in the size hierarchy. These results support the broadly confirmed fact that females feed smaller nestlings more than expected according to their size-hierarchy in the brood (see references above), thus exerting the main control over food allocation. These findings have also been found in other species. For instance, in the hoopoe, females, not only can feed preferentially the smallest nestling, usually placed in the worst position (Ryser et al. 2016), but can also ignore begging calls from small nestlings while forcing to swallow food to a silent larger nestling (Martin-Vivaldi et al. 1999). Another fascinating example of parents having full control over food allocation occurs in magpie (*Pica pica*) nests, especially when parasitized by the great spotted cuckoo (*Clamator glandarius*). Sometimes, both in parasitized and non-parasitized nests, magpie parents, when no nestling is begging, may induce one of them to beg by waking it up by touching it softly with the beak. But surprisingly, in parasitized nests, magpies may ignore cuckoos that beg for food (begging magpies are never ignored) and instead wake up one of their magpie nestlings (coax feedings (Soler et al. 2017)). In the presence of the wooden barriers, the amount of food provided to the smallest experimental nestling by males is lower compared to larger nestlings; in contrast, females fed the smallest nestling not significantly less than the other larger nestlings. This result is in agreement with the “Male manipulation hypothesis”, which predicts that females will provide some food to the smallest nestling to keep it alive to benefit of their intense begging forcing males to work harder (Soler et al. In prep).

In conclusion, our findings show that runts do not starve in asynchronous broods as a consequence of being outcompeted by their larger nestlings, but because both parents ignore their intense begging and prefer to feed one of the larger nestlings.

## Acknowledgements

We are indebted to D. W. Mock and E. Schlicht for their valuable comments on a previous version of this manuscript. We thank Teresa Abaurrea for her assistance with field work.

## Data accessibility

All data that support the findings of this study will be available from the Digibug Digital Repository.

## References

Budden AE, Beissinger SR. 2009. Resource allocation varies with parental sex and brood size in the asynchronously hatching green-rumped parrotlet (*Forpus passerinus*). Behav Ecol Sociobiol. 63:637–647.

Caro SM, Griffin AS, Hinde CA, West SA. 2016. Unpredictable environments lead to the evolution of parental neglect in birds. Nat Commun. 7, 10985.

Cramp S. (ed.) 1985. The Birds of the Western Palearctic. Vol. IV. Oxford: Oxford University Press.

Davis JN, Todd PM, Bullock S. 1999. Environment quality predicts parental provisioning decisions. Proc R Soc B 266: 1791–1797.

Ellis LA, Styrsky JD, Dobbs RC, Thompson CF. 2001. Female condition: a predictor of hatching synchrony in the house wren? Condor 103:587–591.

Engqvist L. 2005. The mistreatment of covariate interaction terms in linear model analyses of behavioural and evolutionary ecology studies. Anim. Behav. 70:4, 967–971.

Gottlander K. 1987. Parental feeding behaviour and sibling competition in the pied flycatcher *Ficedula hypoleuca*. Ornis Scand. 18, 269–276.

Hauber ME, Moskát C. 2008. Shared parental care is costly for nestlings of common cuckoos and their great reed warbler hosts. Behav Ecol. 19:79–86.

Jeon J. 2008. Evolution of parental favoritism among different-aged offspring. Behav Ecol. 19:344–352.

Kilner RM. 1995. When do canary parents respond to nestling signals of need? Proc R Soc Lond B. 260:343–348.

Krebs EA, Cunningham RB, Donnelly CF. 1999. Complex patterns of food allocation in asynchronously hatching broods of crimson rosellas. Anim. Behav. 57:753–763.

Lack D. 1947. The significance of clutch size. Ibis 89:302–352.

Lack D. 1954. The Natural Regulation of Animal Numbers. Oxford: Clarendon Press.

Lahaye SEP, Eens M, Iserbyt A, Groothuis TGG, de Vries B, Muller W, Pinxten R. 2015. Influence of mate preference and laying order on maternal allocation in a monogamous parrot species with extreme hatching asynchrony. Horm Behav. 71:49–59.

Lenth RV. 2016. Least-squares means: the R package lsmeans. J Stat Softw. 69:1–33.

Leonard M, Horn A. 1996. Provisioning rules in tree swallows. Behav Ecol Sociobiol. 38:341–347.

Lifjeld JT, Breiehagen T, Lampe HM. 1992. Pied Flycatchers failed to use nestling size as a cue to favour own genetic offspring in a communally raised brood. Ornis Scand. 23:199–201.

Macnair MR, Parker GA. 1979. Models of parent-offspring conflict. III. Intra-brood conflict. Anim Behav. 26:111–122.

Magrath RD. 1990. Hatching asynchrony in altricial birds. Biol Rev. 65:587–622.

Malacarne G, Cucco M, Bertolo E. 1994. Sibling competition in asynchronously hatched broods of the pallid swift (*Apus pallidus*). Ethol Ecol Evol. 6:293–300.

Martin-Vivaldi M, Palomino JJ, Soler M, Soler JJ. 1999. Determinants of reproductive success in the Hoopoe *Upupa epops*, a hole-nesting non-passerine bird with asynchronous hatching. Bird Study. 46:205–216.

McRae SB. Weatherhead PJ, Montgomerie R. 1993. American robin nestlings compete by jockeying for position. Behav Ecol Sociobiol. 33:101–106.

Mock DW, Dugas MB, Strickler SA. 2011. Honest begging: expanding from signal of need. Behav Ecol. 22:909–917.

Mock DW, Parker GA. 1997. The Evolution of Sibling Rivalry. Oxford: Oxford Univ. Press.

Mock DW, Schwagmeyer PL, Dugas MB. 2009 Parental provisioning and nestling mortality in house sparrows. Anim Behav. 78:677–68.

Nakagawa S, Johnson PCD, Schielzeth H. 2017. The coefficient of determination R 2 and intra-class correlation coefficient from generalized linear mixed-effects models revisited and expanded. J R Soc Interface. 14:20170213.

Nakagawa S, Schielzeth H. 2013. A general and simple method for obtaining R2 from generalized linear mixed-effects models. Methods Ecol Evol. 4:133–142.

Ostreiher R. 2001. The importance of nestling location for obtaining food in open cup-nests. Behav Ecol Sociobiol. 49:340–347.

Parker GA, Royle NJ, Hartley IR. 2002. Begging scrambles with unequal chicks: interactions between need and competitive ability. Ecol Lett. 5:206–215.

Pinheiro J, Bates D, Debroy S, Sarkar D. 2014. nmle: Linear and nonlinear mixed effects models. R package version 3.1-117. Available: http://CRAN.Rproject.org/package=nlme.

Ploger BJ, Medeiros MJ. 2004. Unequal food distribution among great egret *Ardea alba* nestlings: parental choice or sibling aggression? J Avian Biol. 35:399–404.

R Core Team 2019. R: A language and environment for statistical computing. R Foundation for Statistical Computing, Vienna, Austria. URL https://www.R-project.org/.

Ryser S, Guillod N, Bottini C, Arlettaz R, Jacot A. 2016. Sex-specific food provisioning patterns by parents in the asynchronously hatching European hoopoe. Anim. Behav. 117:15–20.

Slagsvold T. 1997a. Brood division in birds in relation to offspring size: sibling rivalry and parental control. Anim Behav. 54:1357–1368.

Slagsvold T. 1997b. Is there a sexual conflict over hatching asynchrony in American robins? Auk 114:593–600.

Smiseth PT, Amundsen T. 2002. Senior and junior nestlings in asynchronous bluethroat broods differ in their effectiveness of begging. Evol Ecol Res. 4:1177–1189.

Smiseth PT, Bu RJ, Eikenaes AK, Amundsen, T. 2003. Food limitation in asynchronous bluethroat broods: effects on food distribution, nestling begging, and parental provisioning rules. Behav Ecol. 6:793–801.

Smith MG, Dickinson JL, Rush A, Wade AL, Yang DS. 2017. Western bluebird parents preferentially feed hungrier nestlings in a design that balances location in the nest. Behav Ecol Sociobiol. 71: 58.

Soler M, Macías-Sánchez E, Martín-Gálvez D, de Neve L. 2017. Complex feeding behaviour by magpies in nests with great spotted cuckoo nestlings. J Avian Biol. 48:1406–1413.

Soler M. 2001. Begging behaviour of nestlings and food delivery by parents: the importance of breeding strategy. Acta Ethol. 4, 59–63.

Stamps J, Clark A, Arrowood P, Kus B. 1985. Parent-offspring conflict in budgerigars. Behaviour 94:1–40.

Stamps J, Clark A, Kus B, Arrowood P. 1987. The effects of parent and offspring gender on food allocation in budgerigars. Behaviour. 101:177–199.

Tanner M, Kölliker M, Richner H. 2008. Differential food allocation by male and female great tit, *Parus major*, parents: are parents or offspring in control? Anim Behav. 75:1563–1569.

